# Wild Florida Mottled Ducks demonstrate strong heterogeneity in their humoral innate immune response

**DOI:** 10.1101/2024.10.12.617993

**Authors:** Andrea J. Ayala, Matthew Cheng, Thomas A. Hellinger, K. Mark McBride, Jonathan Webb, Andrew Fanning, Paul Snyder, Margherita Ferragamo, Samantha C. Garcia, Nyah Sterner, Karyn L. Bischoff, Salvador-Almagro Moreno, C. Brandon Ogbunugafor

## Abstract

The Florida Mottled Duck (*Anas fulvigula fulvigula*) is a unique subspecies of waterfowl whose range is limited to peninsular Florida, USA. As an endemic species, Florida Mottled Ducks face numerous conservation stressors, such as habitat conversion and hybridization with non-native Mallards (*Anas platyrhynchos*). In addition to these numerous stressors, Mottled Ducks are also contending with emerging and/or geographically expanding waterborne pathogens such as *Vibrio* spp., due to the effects of climate change. However, even given their conservation needs, little is known with respect to the health, physiology and the immunity of wild Mottled Ducks in Florida. Given this lack of data, we performed health assessments of Mottled Ducks in the Central Florida area.

Specifically, we examined the humoral innate immune system, i.e., plasma, of Mottled Ducks in response to a common, but extraneous pathogen, *Escherichia coli* strain ATCC number 8739. We utilized a bactericidal assay (BKA) commonly used in eco-immunology, to provide insight into the bactericidal capacities of captured Florida Mottled Ducks. We statistically tested the BKA capacity of 23 Mottled Ducks in response to age and whole blood lead levels (Pb). We found that there was no statistically significant relationship between the covariates we measured and Mottled Duck BKA capacity against *E. coli*. However, the variability we observed in the BKA capacity of this subspecies warrants further research into additional physiological and ecological covariates coupled with potential immune stressors that Florida Mottled Ducks may be contending with.

## Introduction

The Florida Mottled Duck (*Anas fulvigula fulvigula*) is an iconic and endemic subspecies that is contending with immense conservation pressures [1–3]. Urbanization, agricultural conversion, and suburban sprawl continue to fragment the habitat of this species, an observation that was initially reported in the 1950’s [3–5]. Mottled Ducks are resident dabbling ducks, i.e., ducks that ‘tip-up’ in the water to forage, that inhabit a gradient of urban to rural habitats throughout the state [4]. Currently, populations in northern Florida are facing stronger environmental stressors than those in the southern portion of their range [2]. However, the most pressing range-wide management concern remains their hybridization with non-native Mallards (*Anas platyrhynchos*) [1, 3]. Due to the sheer number of Mallards and Mottled Duck x Mallard hybrids in Florida, the genetic swamping of ‘pure’ Mottled Ducks is an overwhelming priority with regards to their preservation as a subspecies [1, 6].

In addition to urbanization and genetic introgression, aquatic species such as Mottled Ducks are now contending with a rise in waterborne pathogen abundances and distributions due to climate change [7, 8]. However, even given their conservation needs, little is known with respect to the health and physiology of wild Mottled Ducks in Florida. Specifically, we were interested in understanding the capacity of Mottled Ducks to kill an invasive bacteria (*Escherichia coli* strain ATCC number 8739) using a sample of their fresh, frozen plasma [9, 10]. Such physiological traits, i.e., immune function, may be linked to their health or fitness [11, 12]. Commonly known as the bactericidal assay (hereafter, BKA), it is a measure of humoral innate immunity – the first line of defense against pathogen invasion [13, 14].

As one of two immunological strategies employed, together, the humoral and the cellular avian innate immune system confers a rapid, initial resistance against circulating pathogens - including novel and emerging infectious diseases [15]. It does so through the use of cellular effector proteins and pattern recognition receptors, which recognize antigenic constituents such as complex polysaccharides, glycolipids, lipoproteins, nucleotides and nucleic acids [16, 17].

Conversely, the adaptive immune system mounts a slower, albeit more sophisticated response against a single antigen which can be recalled from immunological memory cells upon secondary exposure to that specific antigen [18, 19]. Both systems work in tandem to protect the host from microbial invasion, however, the strength of that response can vary according to sex, age, body condition, pace of life and pollutant exposure [20–27].

For instance, lead (Pb) is a highly toxic heavy metal with immunosuppressive properties in vertebrates [28]. In birds, Pb interferes with innate and adaptive cell-mediated, and humoral immunity [27, 29–31]. Although the use of lead shot for hunting waterfowl was banned in 1991 in the United States, followed by Canada in 1997 [32], it persists in terrestrial and aquatic habitats as a legacy contaminant [33, 34]. Fishing tackle is another source of Pb which may be consumed by aquatic birds as well [35, 36]. Waterfowl ingest lead primarily through feeding, with dabbling ducks in the Mallard complex amongst those most affected [37, 38]. In 1986, the U.S. Fish and Wildlife Service mandated that blood Pb levels above 20 ug/dl in waterfowl was a sublethal threshold that exceeded background exposure levels [37, 39].

Crucial to an organism’s survival is the ability to generate energetic trade-offs amid resource limitations and environmental stressors [40–43]. While the maintenance of a robust immune system is critical, so too is the ability of birds to undergo physiologically demanding life stages such as migration, molt, or reproduction [25, 40, 44, 45]. In such instances, immunocompetence may be negatively correlated with such natural history investments. Thus, there are many benefits to utilizing the BKA to measure humoral and/or adaptive innate immunity, especially in a species experiencing numerous conservation stressors [12, 46]. For example, it is relatively easy to perform in a field or laboratory setting, it requires only small volumes of blood, serum or plasma, and the assay’s interpretation is straightforward [47, 48].

The BKA has been widely implemented by eco-immunologists to investigate relevant avian life history variables in relation to immunocompetence [49, 50]. For example, infestations of chewing lice (*Mallophaga* spp.) were associated with a reduction in the microbicidal killing capacity of suburban birds in Chicago, Illinois, USA [14]. Baseline cortisol levels were positively linked to the bactericidal ability of male Red-winged Blackbirds (Agelaius phoeniceus) in Santa Barbara, California, USA [51]. Thermal variation in egg incubation, specifically higher temperatures, covaried with a higher BKA capacity in nestling American Robins (*Turdus migratorius*) [52]. House Finches (*Haemorhous mexicanus*) infected with the bacterial pathogen *Mycoplasma gallisepticum* had a higher BKA capacity in spite of handling stress in contrast to their uninfected counterparts [53]. In essence, such examples are only a limited representation of how the BKA can help us understand the factors that influence avian immunology [49].

Given the pressing conservation needs of Florida Mottled Ducks, we performed BKAs on free-ranging individuals captured in Central Florida to derive insight into the humoral innate immune status of the population, as well as in response to Pb exposure levels and age. We hypothesized that older birds, having had more time to be exposed to immunosuppressive environmental Pb, would have a diminished BKA capacity. Thus, using fresh, frozen plasma that was collected during a pathogen surveillance study, we explored Mottled Duck BKA capacity against *E.coli* ATCC strain 8739. Our statistical analyses measured our BKA results in relation to covariates such as age, sex, weight in grams, and whole blood lead (Pb) levels. Interestingly, we found that while BKA results were highly variable across individuals, there were no clear statistically significant patterns with respect to the surveyed demographic variables or Pb levels.

## Materials and Methods

### Ethics statement

All fieldwork was conducted under U.S. Geological Service Bird Banding Laboratory federal permit number 06672, in collaboration with the Florida Fish and Wildlife Conservation Commission. In addition, all work was conducted in accordance with Yale University’s Institutional Animal Care and Use Committee (IACUC) protocol AUP #: 2021-20379. All Mottled Ducks were captured on public lands owned by the state of Florida, Orange County, or the city of Orlando.

### Capture and sampling of Florida Mottled Ducks

Florida Mottled Ducks (n = 42) were captured at three sites in Central Florida during August of 2022. Capture sites included one suburban site (Lake Millenia, Orange County, n = 1), and two rural sites (Puzzle Lake, Seminole County, n = 3 and T.M. Goodwin Wildlife Management Area, Indian River County, n = 38). Mottled Ducks were captured using either eight-foot wide bownets (Modern Falconry, Eau Claire, WI, USA) that were baited with cracked corn [54], or through night-time spotlighting [55, 56]. All individuals were banded with a federal band as per USGS guidelines [2] and to prevent pseudo-replication [57].

Demographic attributes such as age and sex were assessed using physical characteristics as described by Pyle (2008) [58, 59]. Specifically, sex was assessed using bill color, where males have an olive green to yellow bill, and females have an orange to brown bill, with a distinctive spot on the underside [60]. Age was assessed through plumage molt patterns and cloacal examination [61]. Birds were placed into the hatch-year (HY) category if they had hatched in the 2022 breeding cycle, or after-hatch year (AHY) if hatched before 2022 [61]. The mass in grams for each bird was collected using an MTB 20 medical-grade scale (Adam Equipment, Oxford, CT, USA). Given that measurements were collected during the molting season for Mottled Ducks, including primary wing feathers [62], we were unable to collect wing chord lengths to standardize their size [63].

Up to three mL of blood were collected from the medial metatarsal vein, located on the medial aspect of the tarsometatarsus [64, 65]. This is one of the most commonly utilized sites for blood collection in waterfowl [65]. Up to one mL each was transferred to a red-top vacutainer for serum, a green-top heparinized vacutainer for plasma, and a purple-top vacutainer with EDTA for whole blood lead analysis (Becton Dickinson, Franklin Lakes, NJ, USA). Vacutainers were placed into a cooler with ice packs until transport to the lab. Plasma and serum vacutainers were centrifuged at 5000 RPM for 10 minutes. Whole blood vacutainers were immediately shipped overnight with ice packs to the toxicology department of the Cornell Animal Health Diagnostic Laboratory (AHDC; Cornell University, Ithaca, New York) for analysis [66]. Separated plasma and serum samples were also immediately shipped overnight on ice packs for storage at Yale University and maintained at -20°C until further analyses [67].

### Whole blood lead analysis (Pb)

Whole blood samples in vacutainers with EDTA were prepared for analysis at the AHDC of Cornell University. Briefly, blood samples underwent graphite furnace atomic absorption spectroscopy (GFAAS) Pb analysis using a transversely heated GFAAS with longitudinal Zeeman-effect background correction [68, 69]. A 100-μl subsample was taken of each sample, placed into a 2-ml cup, and mixed with 900 μl of Pb matrix-modifier solution composed of distilled deionized water, 0.02% analytical-grade ammonium phosphate, 0.05% analytical grade magnesium nitrate, 1.0% analytical-grade nitric acid, and 0.1% Triton X. Standards containing 2.50, 5.00, 10.00, and 50.00 μg/dl Pb to construct a calibration curve were prepared by diluting certified atomic absorption standard solution in matrix modifier and placing into 2-ml cups, which were placed in the autosampler tray. All blanks, standards, and samples were analyzed in duplicate. The calibration curve provided a linear response across this range with a correlation coefficient of 0.999. The average of two replicates was taken for statistical analysis. Blood samples containing greater than 50 μl/dl of Pb were diluted 1:1 in matrix-modifier solution and reanalyzed. The limit of detection (LOD) was 2.50 μg/dl based on the lowest standard solution [70].

### Bactericidal assays

A wide-ranging literature search indicated that *E. coli* ATCC strain number 8739 is among the most widely used strains in avian eco-immunology [25, 47, 51, 53, 71–76]. Thus, we incorporated the same strain into this experiment for reproducibility purposes. We prevented cross-contamination by performing all microbial work in an A2 Class II Biosafety Cabinet (Baker, Sanford, ME, USA) and applying sterile principles. Clear plate sealers were used when the 96-well plate was removed from the biosafety cabinet. In addition, all materials such as media, pipette tips, and reagent boats were autoclaved prior to use. Bactericidal assays were performed using a protocol delineated by French and Neuman-Lee [77] with minor modifications as described in LaVere et al [78]. Any deviations from these protocols are detailed below.

Preparations to perform the BKA are briefly described as follows. Fresh tryptic soy broth (Sigma-Aldrich NO. T8907) was constituted by dissolving 15 grams in 500 mL ultrapure filtered water (Millipore Milli-Q™ Direct Water Purification System, Sigma-Aldrich). Fresh tryptic soy agar petri dishes (Sigma-Aldrich) were made by dissolving 22.5 g in 500 mL ultrapure water and storing them at 4°C. We then incubated 100 mL of 0.9% phosphate buffered saline (PBS) at 37°C and added the lyophilized *E. coli* pellet (EPower Microorganisms, Microbiologics).

Finally, the tryptic soy agar plates were inoculated with the stock solution of *E. coli*, and incubated for 12 hours at 37°C. To perform the BKA, the Prompt Inoculation System (BD BBL™ Prompt™ Inoculation System, Franklin Lakes, NJ, USA) [79] was used to create a 10^5^ working stock of *E. coli* from five colonies grown on tryptic soy agar [78]. The Mottled Duck frozen plasma samples (n = 27) were placed on ice and thawed completely. Flat-bottomed, 96 well plates were used to run the assay which included positive controls (125 uL of tryptic soy broth + 20 uL of PBS + 4 uL of *E. coli*), negative controls (125 uL of tryptic soy broth + 24 uL of PBS), sample wells (125 uL of tryptic soy broth + 16 uL of PBS + 4 uL of E. coli + 4 uL of plasma) and blank wells (125 uL of tryptic soy broth). The optimal dilution of plasma for the BKA was found to be 1:2 (equal parts plasma and PBS) after testing a range of concentrations. PBS, plasma, and *E. coli* were added to wells first, then shaken using a microplate shaker at 300 RPMs for one minute to ensure proper mixing. Following this, tryptic soy broth was added to the appropriate wells and shaken again.

Plasma samples were run in triplicate, with the exception of two samples. One sample had six replicates, and one sample only had only a single replicate due to limited plasma.

Positive, negative, and blank wells were each assayed in replicates of eight. An accuSkan FC (Fisherbrand, Waltham, MA, USA) microplate reader was used to analyze the optical density of each well at 405 nm [80]. After the initial reading, the plate was incubated at 37°C for 12 hours, after which the plate was analyzed again. Finally, the BKA capacity for each plasma sample was calculated as per LaVere [78].

### Statistical analyses

All statistical analyses were performed using the R platform, version 4.2.2 [81].

#### Statistical analyses for lead (Pb)

Due to standard limitations, the limit of detection (LOD) of all blood lead concentrations were censored at values below 2.50 μg/dl [70]. In light of the already limited sample size with which to test the effects of Pb, we therefore imputed the Pb values that were below the LOD into our dataset [82]. Using a relatively new published method by Hebers et al. (2021), we utilized the lognormal distribution and the probability distribution function of our known values to calculate the unknown values [83]. We then used the mean of the newly estimated values for those Pb values below 2.50 μg/dl and imputed that mean into our dataset [82].

As Pb served as a continuous variable, we assessed its distribution using the Cullen and Frey distribution plot [84], followed by a Shapiro-Wilk’s test of Normality [85].

#### Statistical analysis for the BKA

First, the mean for each set of triplicate plasma samples was calculated. Given that it served as a continuous variable, we also then assessed its distribution using the Cullen and Frey distribution plot [84], followed by a Shapiro-Wilk’s test of Normality [85].

#### Statistical analysis for weight in grams

Also serving as a continuous variable, we assessed its distribution using the Cullen and Frey distribution plot [84], followed by a Shapiro-Wilk’s test of Normality [85].

#### Statistical analyses for age and sex

Each of these is a categorical variable. Age was divided into two categories: HY or hatching-year juveniles, while adults were labeled as AHY, or after hatching year birds. Based on plumage and bill characteristics, Mottled Ducks were also placed into either the male (M) or female (F) category. Both of these classifications are part of the logistic distribution; thus, no Cullen and Frey distribution plot was performed.

#### Bivariate tests of association

Each independent variable was assessed with respect to one another to reduce the likelihood of multicollinearity [86] in our generalized linear model. Due to the non-parametric nature of the variable Pb and Weight in Grams, we performed a Spearman’s Rank Correlation test to examine the association between the two variables. Pb and Age, and Pb and Sex were both analyzed using the non-parametric Mann-U Whitney test. Weight in Grams is a parametric variable, and thus was assessed using an independent samples t-test against Age and Sex, respectively.

#### Generalized linear model of covariates

We assessed the variables Pb and Age against the dependent variable BKA using a generalized linear model (GLM). We did so to test the hypothesis that older birds, having had a longer period by which to be exposed to immunosuppressive environmental lead sources, would have a reduced BKA capacity. We used a GLM given that our independent and dependent variables did not meet the parametric assumptions required to perform an ANCOVA. We further examined the effect sizes of Pb and Age on BKA capacity using the Eta^2^ test.

## Results

### Whole blood lead analysis (Pb)

Pb served as an independent variable. In total, we collected sufficient blood to test 36 (n = 36) of the 42 captured Mottled Ducks for Pb analysis. Eighteen of those 36 individuals had blood Pb levels below the 2.50 μg/dl LOD. Based on the probability distribution function of the lognormal distribution, we calculated a mean value of 1.2 μg/dl, which we imputed for each bird below the LOD.

With respect to our dependent variable, BKA, we had 27 (n = 27) Pb samples for statistical comparison. However, four individuals were unable to be analyzed due to a lack of corresponding data covariates: Pb testing or weight in grams. Thus, in this group of (n = 23) Mottled Ducks, Pb values ranged from a minimum of 1.2 μg/dl to a maximum of 29.46 μg/dl. The overall mean Pb value was 6.03 μg/dl with a standard deviation of 7.05 μg/dl, and a median value of 2.51 μg/dl. The Shapiro-Wilk’s test of Normality found that the Pb data were not normally distributed (W = 0.72952, p-value = <0.0001). According to the Cullen and Frey plot, the variable Pb fell into the Beta distribution, with an estimated skewness of 2.037587 and an estimated kurtosis of 7.741181.

### Weight in grams

The variable weight in grams served as an independent covariate, from which we sub-sampled 23 (n = 23) individuals that corresponded to the birds with plasma samples. In this sub-sample, Mottled Duck weight in grams ranged from a minimum of 460 grams to a maximum of 1200 grams. The mean weight was 907 grams, with an estimated standard deviation of 171.3 and a median of 915 grams. The Shapiro-Wilk’s test of Normality established that this variable was normally distributed (W = 0.96722, p-value = 0.6227). According to the Cullen and Frey plot, the estimated skewness was -0.565771, while the estimated kurtosis was 3.657986.

### Bactericidal assay

BKA served as the sole dependent variable. Corresponding plasma was available for 23 (n = 23) of the 42 captured Mottled Ducks for use in the assay. BKA values ranged from a minimum of 0.1416547 (the maximum killing capacity) to a maximum of 1.143873 (the lowest killing capacity). The mean killing capacity was represented by the value 0.7192972, with a standard deviation of 0.3291034 and a median of 0.6910914. The Shapiro-Wilk’s test of Normality found that the BKA data were not normally distributed (W = 0.90455, p-value = 0.03141). According to the Cullen and Frey plot, the distribution of the BKA values fell between the Beta and Uniform distributions, with an estimated skewness of -0.1391962, and an estimated kurtosis of 1.575785.

### Age and sex

From our sub-sample of 23 (n = 23) Mottled Ducks, 18 birds were AHY and five were HY birds. In addition, eight birds were female, and 15 birds were male.

### Bivariate tests of association

#### Lead and Weight in Grams

Due to the non-parametric nature of the variable Pb, we performed a Spearman’s Rank Correlation test to examine the association between the two variables. We found that the relationship between the two variables was non-significant (*S* = 1794, *p*-value = 0.6056, *ρ* = 0.1136471).

### Lead and Age

A non-parametric Mann-U Whitney test was conducted to determine the statistical significance between Pb and Age. We found that there was no statistically significant relationship between the two variables (*W =* 50*, p*-value = 0.7224). However, the boxplot (Figure 1), suggests that AHY birds have a higher median exposure to Pb than HY birds.

**Figure 1.**
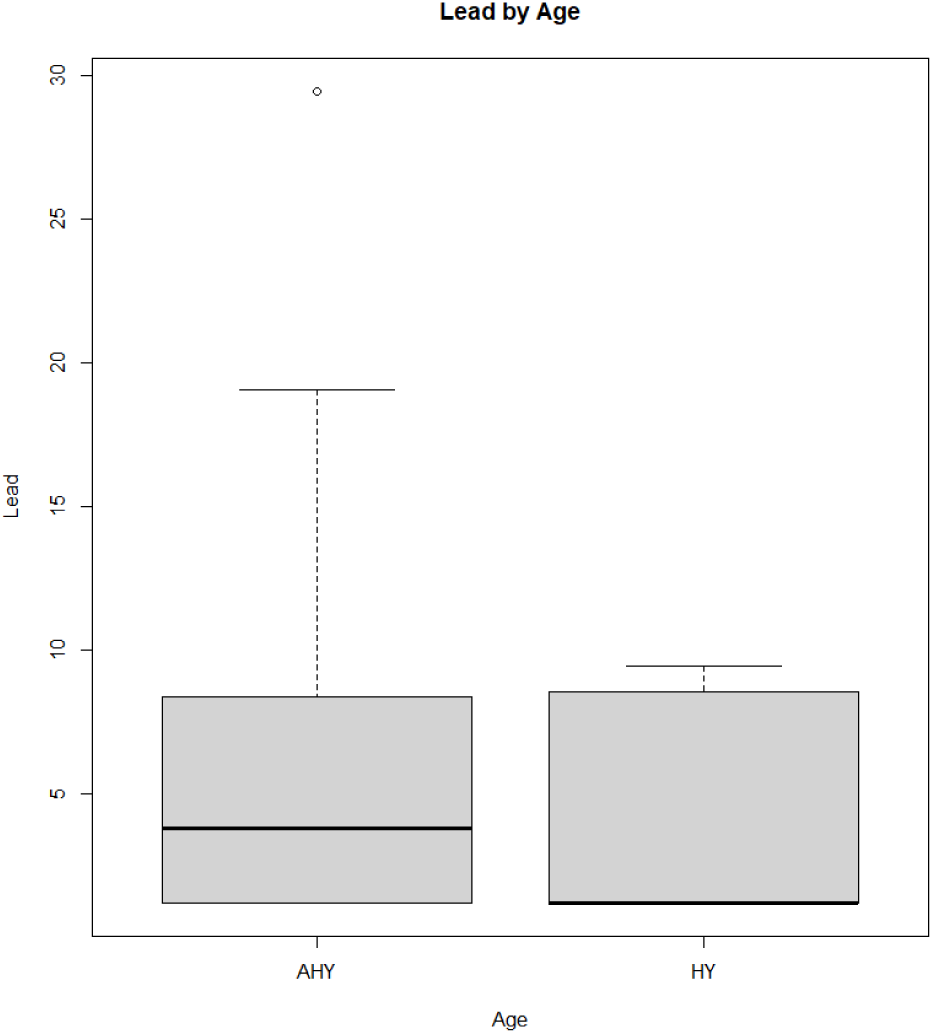
The boxplot above demonstrates the median differences in whole blood Pb levels at the time of sampling between AHY (older) birds and HY (younger) birds. This result was not statistically significant, however (W = 50, p-value = 0.7224).

#### Lead and sex

A non-parametric Mann-U Whitney test was also conducted to determine the statistical significance between Pb and Sex. We found that there was no statistically significant relationship between the two variables (*W* = 56*, p*-value = 0.8109). However, the boxplot (Figure 2), suggests that males have a higher median exposure to Pb than female birds.

**Figure 2.**
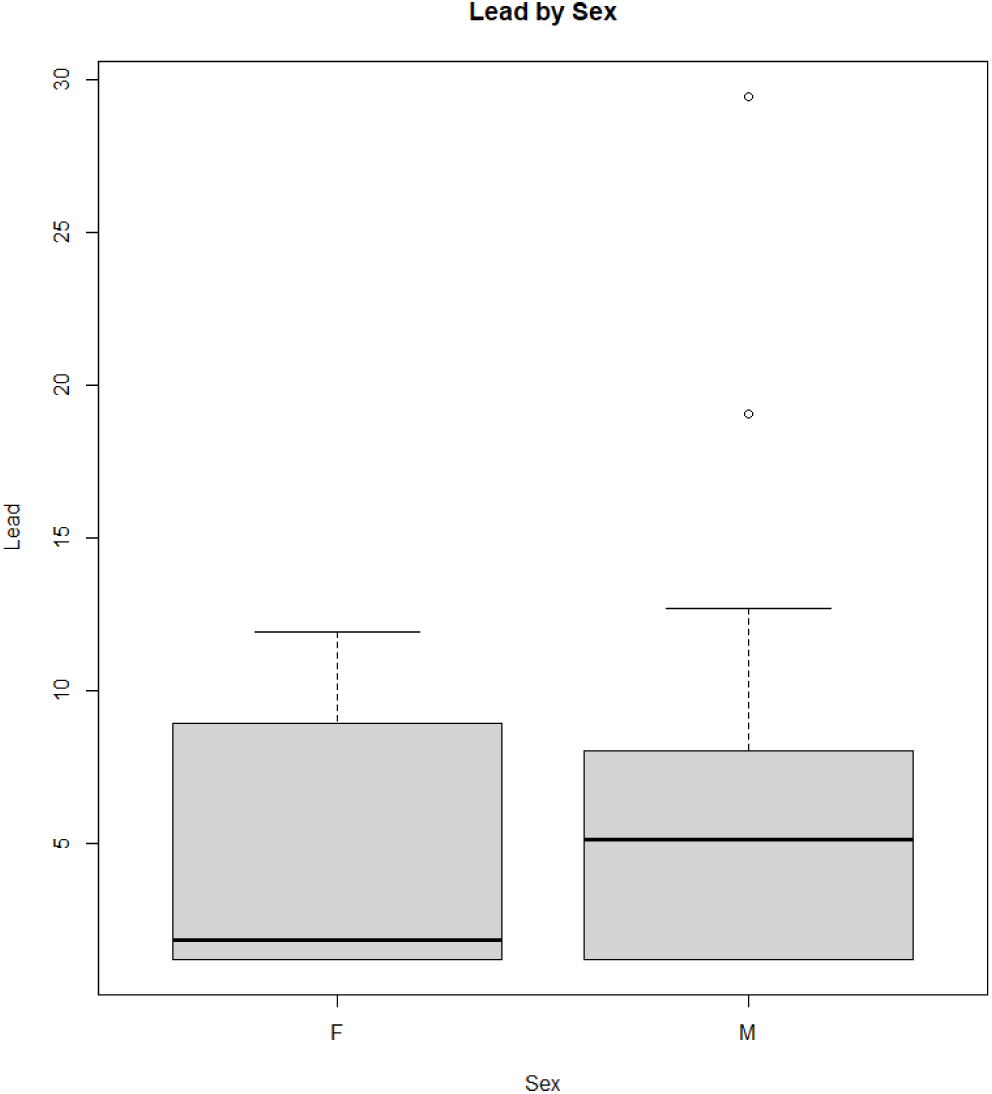
The boxplot above demonstrates the median differences in whole blood Pb levels at the time of sampling between F (female) birds and M (male) birds. This result was not statistically significant, however (W = 56, p-value = 0.8109).

### Weight in grams and age

We performed a parametric independent samples t-test between the variables weight in grams and age for our sub-sample of n = 23 birds. Although weight was not statistically significant (*t* = 1.725, *df* = 21, *p*-value = 0.09922) when contrasted between the groups AHY and HY, there was a distinct trend towards older birds categorized in a higher weight class. AHY birds had a mean weight in grams of 938 g and HY birds had a mean weight of 795 g.

### Weight in Grams and Sex

We again performed a parametric independent samples t-test between the variables weight in grams and sex for our sub-sample of n = 23 birds. In this instance, weight in grams was statistically significant (*t* = -4.3335, *df* = 21, *p*-value = 0.0002927) when contrasted between the groups F and M. Male birds had a greater mean weight in grams of 991 g and F birds had a mean weight of 749 g.

### Generalized linear model

In our GLM, we examined the effects of Pb and Age with respect to BKA capacity as our dependent variable. A non-significant effect was observed for Pb (*t* = -0.243, *df* = 20, *p*-value = 0.810, η² = 0.0002) and Age (*t* = -1.368, df = 20, *p*-value = 0.187, η² = 0.09) on our BKA values. Below, the histogram demonstrates the variability of the BKA capacity in our 23 Mottled Duck samples (Figure 3).

**Figure 3.**
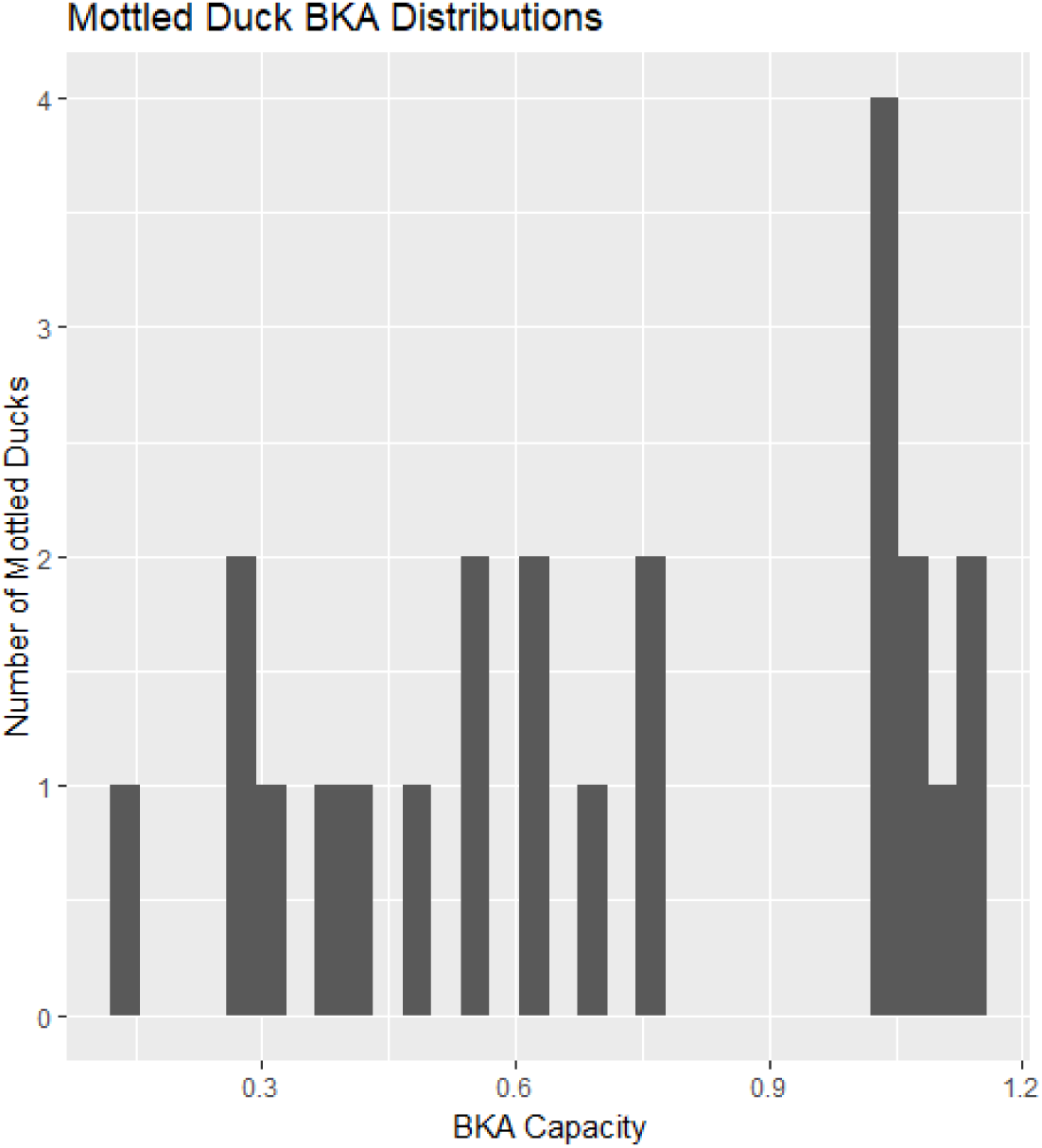
The histogram demonstrates the BKA capacity of our Mottled Ducks across a sample size of 23 birds, both HY and AHY, as well as M and F. Greater BKA capacity is represented by a lower value along the x-axis, while a higher value along the x-axis is indicative of diminished BKA capacity.

## Discussion

A 2019 study stated that the “variation in immune defense influences infectious disease dynamics within and among species” [49]. In this study, we sought to characterize the BKA capacity of Mottled Ducks in Central Florida as a proxy for the function of their humoral innate immune system [22, 87]. Given the array of stressors that Mottled Ducks are currently facing in Florida [1, 2, 56, 59, 60], we found it was imperative to quantify their innate immune resiliency in response to an additional pressure: invasion by novel and/or emerging pathogens due to climate change, i.e., *Vibrio* spp. [88, 89]. We found that the microbial killing capacity of Mottled Ducks varied widely among individuals (Figure 3), however we were unable to decipher a specific covariate that we had measured to explain why.

Although our limited sample size of 23 Mottled Ducks presents difficulties in making further inferences [90–92], we were surprised that birds with higher Pb levels did not have a statistically significantly reduced BKA capacity [36, 93–95]. However, a trade-off may be in place that we had not initially considered. Specifically, while AHY birds may have had longer environmental contaminant exposures to Pb, they also generally have a much more robust immune system than HY birds [96, 97]. HY birds are also more sensitive to the immunotoxic effects of Pb [31]. While the relationship between age and immunity has been documented more frequently in the case of the humoral adaptive immune system [98–100], the innate immune system may be an important trait that covaries with age as well [23, 101, 102]. More research is needed into the interplay of age, environmental contaminants, and innate immunity among avian species.

The expenditure of energetic resources during the summer or remigial molt [59, 103–105], and the subsequent differences in those molt patterns that are delineated by age may have played a role in the BKA value variation we observed [97, 106, 107]. For example, Rufous-collared Sparrows (*Zonotrichia capensis peruviensis*) in Chile that were not molting and not breeding had the lowest levels of immune function as measured by the BKA [44]. However, this pattern should have been apparent between molting HY and AHY Mottled Ducks given differing molt patterns [59, 62, 108], and yet that trend was not statistically significant according to our analyses.

We are able to report a notable finding: there are significant differences in the weight in grams between males and females, however these weights were not standardized by wing chord or tarsus length [109]. Further research into the Mottled Duck system will require a larger sample size of birds captured across multiple seasons to strongly infer the covariates that may be influencing their health in response to the stressors they are facing.

This study sought to explore the physiological consequences of the challenges that Mottled Ducks are facing in an increasingly complex world defined by stressors of various kinds. Future work can further interrogate the interaction between ecological factors and aspects of avian health. The benefits of this research could transcend any particular taxa and speak to broader questions surrounding the biological and health consequences of climate change and anthropogenic changes.

## Data availability

Data and code can be found on Github: https://github.com/OgPlexus/BKA1

## Acknowledgements

The authors would like to thank members of the Almagro-Moreno and Ogbunu research groups for support on various aspects of the project. AJA would like to acknowledge the support of an NSF Postdoctoral Fellowship in Biology, award number: 2010904. AJA would also like to acknowledge funding from the Blake Nuttall Ornithological Society and National Geographic.

